# TENSCell: A 3D-Printed Device for High-Magnification Imaging of Stretch-Activated Cells Reveals Divergent Nuclear Behavior to Different Levels of Strain

**DOI:** 10.1101/742130

**Authors:** Benjamin Seelbinder, Adrienne K. Scott, Isabel Nelson, Stephanie E. Schneider, Kristin Calahan, Corey P. Neu

## Abstract

Mechanical cues from the environment influence cell behavior. Mechanisms of cellular mechanosensation are unclear, partially due to a lack of methods that can reveal dynamic processes. Here, we present a new concept for a low-cost, 3D-printed *TENSCell* (TENSion in Cells) device, that enables high-magnification imaging of cells during stretch. Using this device, we observed that nuclei of mouse embryonic skin fibroblasts underwent rapid and divergent responses, characterized by nuclear area expansion during 5% strain, but nuclear area shrinkage during 20% strain. Only responses to low strain were dependent on calcium signaling, while actin inhibition abrogated all nuclear responses and increased nuclear strain transfer and DNA damage. Imaging of actin dynamics during stretch revealed similar divergent trends, with F-actin shifting away from (5% strain) or towards (20% strain) the nuclear periphery. Our findings emphasize the importance of simultaneous stimulation and data acquisition to capture rapid mechanosensitive processes and suggest that mechanical confinement of nuclei through actin may be a protective mechanism during high strain loads.

**STATEMENT OF SIGNIFICANCE:** Cells can sense and respond to mechanical cues in their environment. These responses can be rapid, on the time scale of seconds, and new methods are required for their acquisition and study. We introduce a new concept for a 3D-printed cell-stretch device that allows for simultaneous high-resolution imaging, while also being low-cost and easy to assemble to enable broad applicability. Using this device, we further demonstrated to importance of simultaneous stimulation and data acquisition to elicit mechanosensitive cell behavior as we observed rapid changes in nuclear size and reorganization of actin filaments around the nuclear border in skin cells. Overall, our results suggest that the rapid reorganization of actin during high loads might protect the genome from strain-induced damage.

## INTRODUCTION

Mechanical cues from the environment are known to have a profound impact on cell fate (1) and cell behavior (2, 3), a phenomenon referred to as mechanosensation. Changes in mechanical properties due to trauma (4, 5), chronic conditions (6–8), or genetic predispositions (9, 10) lead to cellular degeneration and result in a range of pathologies (11). Understanding the mechanisms involved in mechanosensation would help to work towards mitigating these conditions and might allow researchers to direct cell differentiation to generate artificial tissues for drug testing (12) or organ repair (13).

The nucleus is thought to be an essential mechanosensitive organelle as it is tightly connected to all parts of the cytoskeleton through LINC (Linker of Nucleo- and Cytoskeleton) complexes (14, 15). Cyclic stretch has been shown to change nuclear morphology (16–18), induce chromatin condensation (17–19), increase mechanical resistance of nuclei (16, 18, 20) and alter gene expression (21, 22). Disruption of LINC complexes have shown to inhibit stretch-induced changes in chromatin remodeling (23–25) and gene expression (21, 22), suggesting that strain transfer from the cytoskeleton plays an important role for nuclear mechanosensation. The actin skeleton, in particular, has been shown to be crucial for nuclear responses to dynamic mechanical stimulation (18, 21, 23, 26). Despite these findings, the underlying mechanisms of cellular and nuclear mechanosensation pathways are mostly unclear, partially due to a lack of accessible methods to image cell behavior under mechanical stimulation in real time.

Nuclear responses to mechanical stimulation can be very rapid (i.e., on the time scale of seconds) (17, 21) highlighting the importance of simultaneous stimulation and data acquisition to understand these processes. However, commercially available cell stretch devices (e.g. Flexcell) do not allow for the use of high magnification objectives necessary to elucidate single cell behavior (27). Custom-built devices designed by research groups either lack high-resolution live imaging capability (28, 29) or use expensive components such as precise linear actuators (30–32) or optical traps (33). In addition, designs are often complicated and require special expertise, which overall makes them difficult to replicate for widespread use.

Here we propose the design of a low-cost TENSCell (TENSion in Cells) device that uses an electromagnetic piston to apply equiaxial stretch to a thin silicone membrane suspended over a water-immersion objective (**Fig. 1a, Fig. S1**). We used electromagnetic force as it can be precisely controlled without requiring special components, in our case only needing an electromagnetic coil made in-house and a permanent rare earth magnet attached to the piston. Most device parts were 3D printed making the system easy to replicate at a low cost. Only one part, a tapered circular ring that interfaces with the silicone membrane, was machined due to low-friction material requirements. The remaining parts (rare earth magnet, linear rails with carriages and electronics for control) were generally inexpensive and readily available (**Table S1**). Using the TENSCell device, we investigated the dynamic response of nuclei from mouse embryonic skin fibroblasts when exposed to cyclic stretch. Our experiments verified that nuclei respond rapidly to mechanical stimulation. To our surprise, we observed opposing nuclear responses during low (5%) or high (20%) strain regiments, as we observed nuclear area shrinkage and chromatin condensation in response to high strain, but nuclear area expansion and chromatin decompaction in response to low strain. This dichotomous behavior appeared to be mediated through different pathways as only responses to low cyclic strain were dependent on calcium signaling while the actin skeleton was necessary for any strain response. Further investigation of actin dynamics during cyclic stretch revealed a similar divergent behavior with actin filaments shifting towards the nuclear periphery during high loads and towards the cell border during low loads. Actin depolymerization lead to an increase in nuclear strain transfer and DNA damage, overall suggesting that F-actin-dependent nuclear shrinkage might serve as a protective mechanism during high strain loads.

**Fig. 1.**
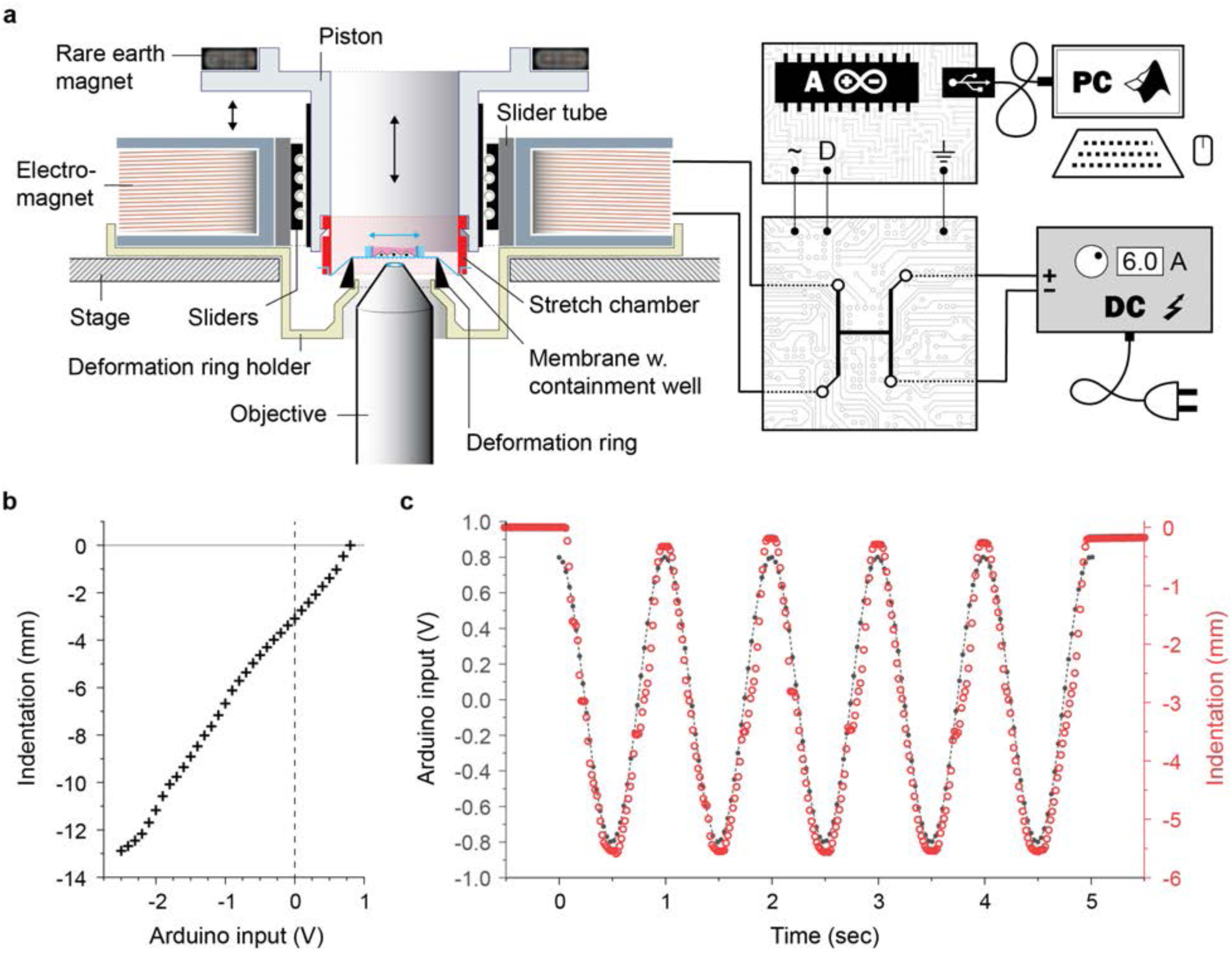
TENSCell device allows for precise and repeatable membrane stretch via electromagnetic force. **a)** A cross-sectional illustration of the stretch device and its control circuit is shown. A suspended piston containing a permanent magnet moves vertically through an electromagnetic coil. Downward motion of the piston stretches a silicon membrane over a deformation ring which holds the stretched membrane at a constant distance over a microscope objective. See Fig. S1b for real images of the device. To control the electromagnet, an H-bridge is used to modulate intensity and direction of a constant current (6 A) from a DC power source through low voltage signals using an Arduino microprocessor. On the Arduino, a PWM pin (∼, power-wave-modulation, 0-5 V) is used to control the intensity and a digital pin (D, 0/1) is used to control the direction of the current. A USB interface enables control of the Arduino inputs via MATLAB. **b)** A distance measurement laser was used to investigate piston movement, and thereby membrane indentation, in response to electromagnetic force as represented by Arduino input voltages. Electromagnetic force could be used for the precise membrane stretch. **c)** A sigmoidal function was programmed in MATLAB to generate Arduino inputs from +0.8 to −0.8 V at a frequency of 1 Hz and piston indentation was recorded over 5 cycles. Electromagnetic force could be used for precise and repeatable membrane indentation.

## RESULTS

### Cell Stretch Device – Concept and Calibration

To record the dynamic responses of single cells at high magnification during mechanical stimulation, we built a system in which a thin (∼127 µm) silicone membrane is stretched over a microscope objective using an electromagnetic piston (**Fig. 1a, Fig. S1a**). A static deformation ring around the objective allowed for a stable imaging plane in *x*, *y* and *z*-directions. The silicone membrane was assembled into a stretch chamber (**Fig. S1b**) and fused to a compliant silicone ring to form a containment well, e.g. to culture cells. The stretch chamber, in turn, could be engaged into a piston that could move freely up and down in an electromagnetic coil via sliders. The piston contained a rare earth magnet at the top to transfer force from the electromagnet below. When current was applied to the coil, the resulting electromagnetic field lifted the piston up or pushed it down depending on the orientation of the magnetic field. The electromagnetic field, in turn, was controlled through an Arduino microprocessor that modulated intensity and the direction of current from a constant DC power source (**Fig. S2a**). A laser displacement sensor was used to show that the electromagnetic field could be used to precisely (**Fig. 1b**) and repeatedly (**Fig. 1c**) control the position of the piston and, therefore, the indentation of the membrane.

To calibrate the TENSCell device, membranes were coated with fluorescent beads. During membrane stretch, bead displacements were recorded on a microscope to calculate the resulting strain using traction force microscopy (34) (**Fig. 2a**). Since the piston has a weight that would stretch the membrane in the absence of a magnetic field, the baseline Arduino input voltage that produced an electromagnetic field to keep the piston suspended without membrane indentation was determined to be +0.8 V (**Fig. S2b**). For calibration, we recorded bead displacements in response to different Arduino inputs from +0.8 to −3.25 V. The resulting current vs. strain calibration curve fit best to a 3^rd^-order polynomial function due to a mild inflection around 0 V; however, a 2^nd^-order polynomial function fit almost equally well and was chosen for simplicity (**Fig. 2b**). Plotting of membrane indentation vs. strain from associated Arduino inputs showed a linear relationship as would be expected from a linear elastic material such as silicone (**Fig. 2c**). Analysis of repeated membrane stretch from +0.8 to −0.9 V, corresponding to 10% strain, showed reliable strain application within 0.5% with no indication of declining or increasing trends within 11 repeats (**Fig. 2d**). Next, we performed live cell imaging of cells during cyclic stretch routines to validate the utility of our device.

**Fig. 2.**
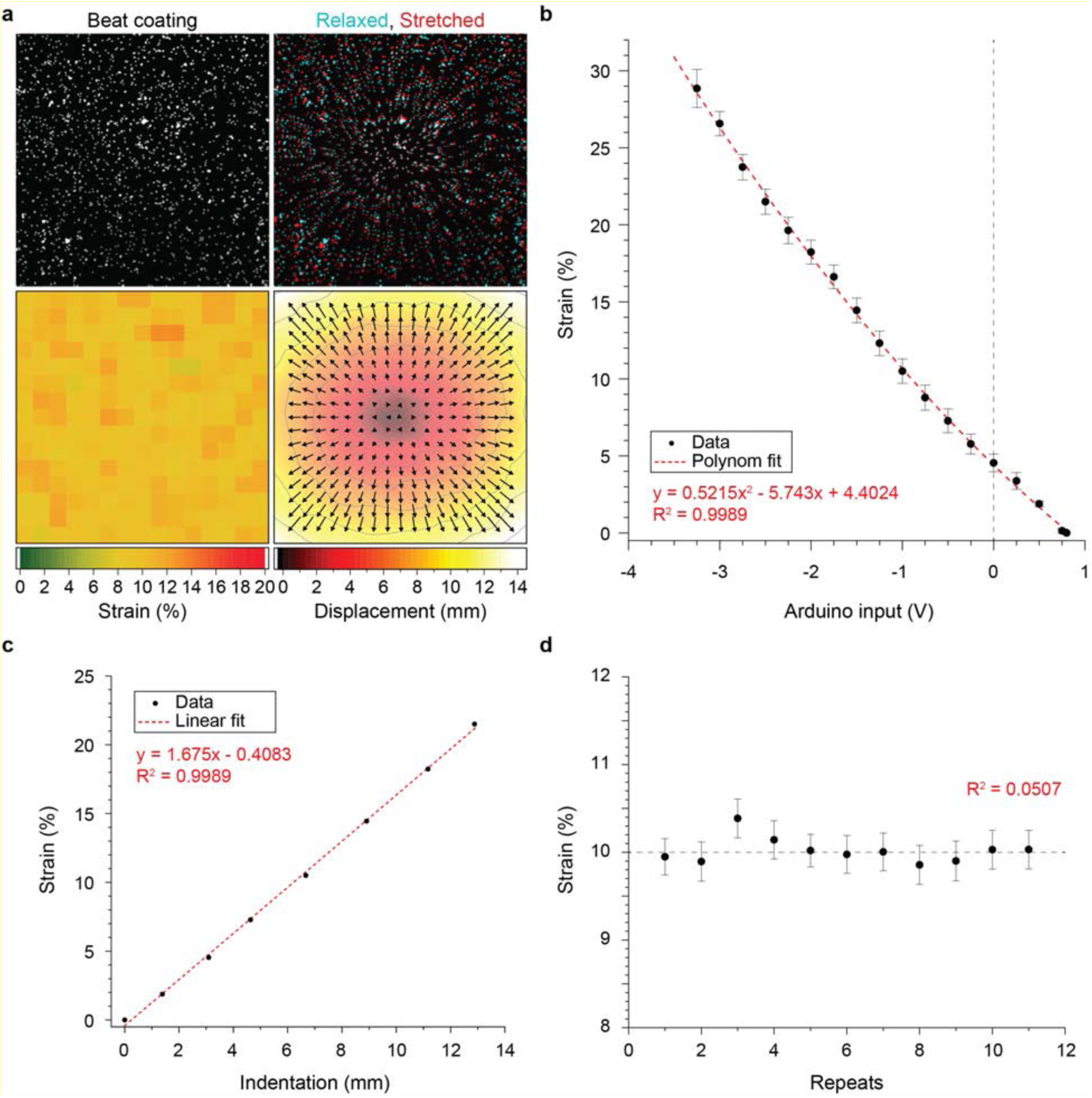
TENSCell device calibration and performance using particle tracking. Membrane containment wells were coated with 2 µm sized fluorescent beads. Images were acquired at baseline (relaxed, Arduino input=+0.8 V, see Fig. S2b) and during stretch to calculate membrane strain from bead displacements, using particle tracking, in response to different Arduino input voltages. **a**) Example of recorded bead displacements and resulting strain map for an Arduino input of −0.9 V. **b)** Calibration curve as determined through bead displacements. The acquired data fit a 2^nd^-order polynomial function; SD; n=3. **c**) Plotting calculated membrane strains over measured membrane indentations from Fig. 1b showed a linear relationship. **d**) Measurements of consecutive membrane indentation for an Arduino input of −0.9 V showed repeatable application of strain within 0.5%.

### Nuclei of MSF Show Dichotomous Responses to Low and High Strain Levels

To test the device, we investigated the dynamic behavior of cell nuclei from mouse embryonic skin fibroblasts (MSF) from H2b-eGFP harboring mice in response to equiaxial stretch. MSFs were seeded onto fibronectin-coated stretch chambers and stimulated with a sigmoidal stretch routine (1 Hz) for 30 min with peak strains of 5%, 10% or 20%. The stretch routine was followed by 30 min of no stimulation (rest) to determine reversibility of any observed responses. In addition to unstretched controls (0%), cells were also subjected to a magnetic field that corresponded to a 20% stretch routine (MAG) under static conditions (no stretch) to assess effects of the magnetic field alone. Image stacks of nuclei were acquired via H2b-eGFP (**Fig. 3a, Movie 1-4**). Unstretched control cells (0%) showed a continuous but slight decline in nuclear area during the 1 h experiment (**Fig. 3b**). In contrast, area of nuclei decreased rapidly (< 2 min) in response to high strains of 20%, continued decreasing during the 30 min of stretch and increased again during the 30 min rest period. Nuclei subjected to 10% cyclic strain also showed a decrease in area but to a lesser extent and more delayed compared to the 20% strain routine. Surprisingly, at 5% strain, nuclear areas increased during stretch and stayed elevated during rest.

**Fig. 3.**
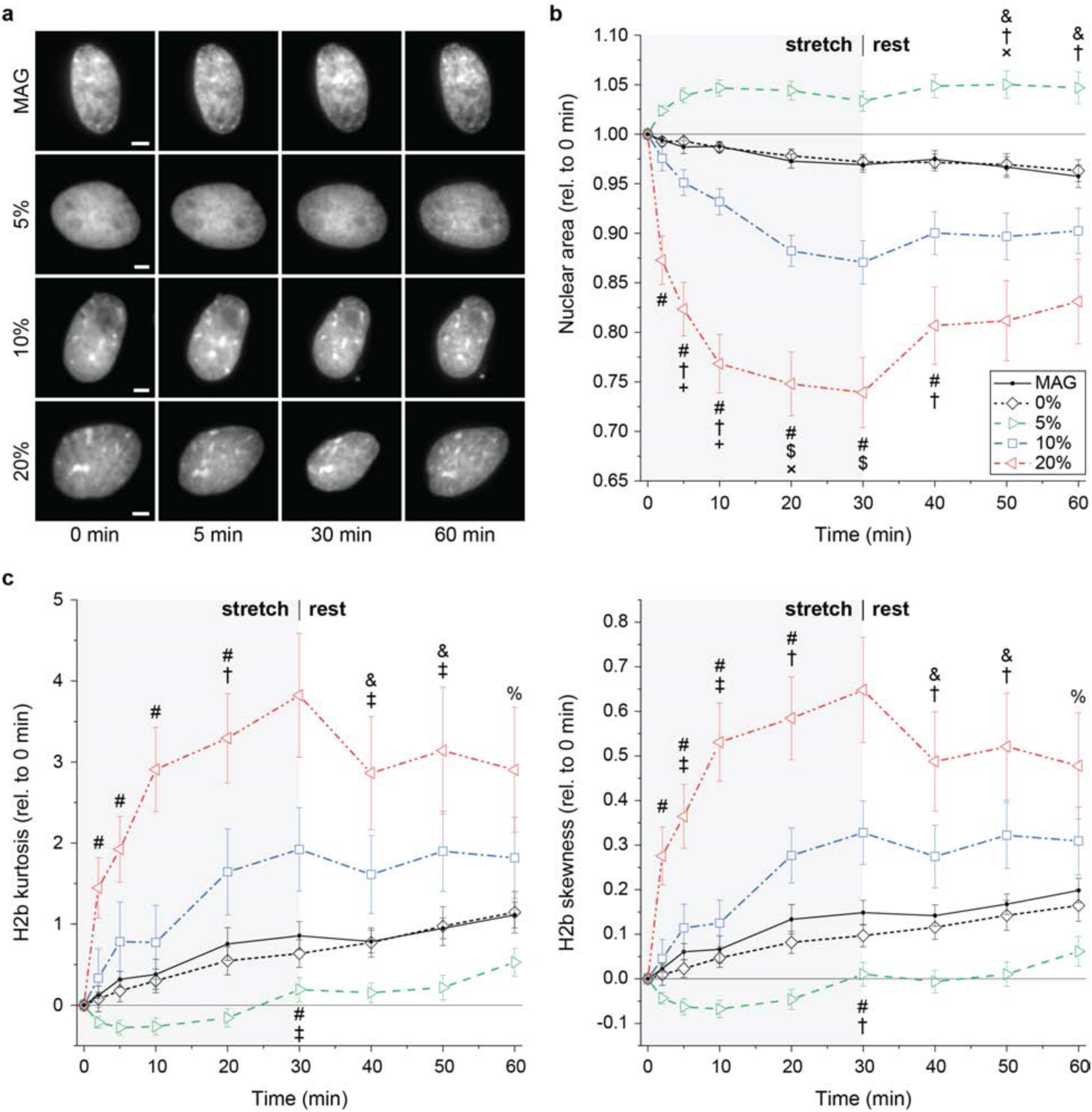
MSF nuclei show opposing changes in nuclear area and chromatin condensation during low strain and high strain cyclic stretch. Mouse embryonic skin fibroblasts were exposed to 30 min of sigmoidal stretch with peak strains of 0%, 5%, 10% or 20%, followed by 30 min of no stimulation (rest), during which image stacks of nuclei were recorded. Control cells were exposed to the magnetic field alone without stretch (MAG). **a**) Images of nuclei recorded via H2b-eGFP; scales=5 µm. **b**) Relative changes in nuclear area (relative to 0 min) during stretch routines. Nuclear areas decreased in response to high strains (10%, 20%), but increased for low strain (5%), while exposure to the magnetic field alone (MAG) showed no difference compared to unstretched cells (0%). **c**) Difference in H2b histogram kurtosis and skewness (compared to 0 min) during stretch routines. Kurtosis and skewness increased under high strain routines (10%, 20%) while they decreased for low strain (5%), indicating elevated or subsided chromatin condensation, respectively; SEM; n>24 from 4 exp.; ANOVA: # (p<0.01) for 20% vs. all, $ (p<0.01) for 10% vs. all, † (p<0.01) or ‡ (p<0.05) for 5% vs. 10%, & (p<0.01) or % (p<0.05) for 20% vs. MAG, 0% and 5%, + (p<0.05) for 5% vs. MAG, × (p<0.05) for 5% vs. MAG and 0%.

We further analyzed chromatin condensation as measured by the changes in skewness and kurtosis of chromatin intensity histograms, with a shift towards higher intensities (positive skewness) and narrowing of the histogram peak (positive kurtosis) indicating elevated chromatin compaction (35). Similar to nuclear area, MSF nuclei subjected to 20% cyclic strain showed a rapid increase in both skewness and kurtosis during the stretch interval, followed by a mild decline during the rest period. Nuclei subjected to 5% cyclic strain showed a decrease in skewness and kurtosis during the stretch and rest period (**Fig. 3c**). Changes in nuclear area as well as H2b histogram skewness and kurtosis of MSFs subjected to the magnetic field alone (MAG) closely matched that of unstretched control cells (0%) in absence of a magnetic field, suggesting that the magnetic field had no influence on the observed cell behavior.

To test whether chromatin dyes are suitable to investigate mechanosensitive behavior of nuclei, MSFs were subjected to a 20% strain routine after staining with Hoechst (**Fig. S3**). Stained nuclei showed a similar dynamic of area decline during stretch, but no recovery during the rest period, compared to unstained nuclei. Additionally, measures of chromatin condensation were reduced during stretch, overall suggesting that the use of chromatin dyes can alter nuclear responses to stretch. Together, these results further confirmed that nuclei can respond rapidly to mechanical stimulation, which emphasizes the need for live imaging capabilities to capture these effects. Moreover, we were surprised to find that MSFs nuclei showed divergent responses to different levels of strain, as nuclear areas were increased and chromatin compaction decreased for low strains and *vice versa* for high strains.

### Nuclear Responses of MSFs to Low, but Not High Cyclic Strain, are Sensitive to Calcium While Actin is Essential for Both

We observed a dichotomous change in nuclear area and chromatin condensation of MSF nuclei in response to high and low cyclic strain. Next, we aimed to investigate whether these responses were mediated through different signaling pathways. Stretch-induced chromatin condensation has been shown to be dependent calcium signaling (17). Treatment of MSFs with BAPTA (BP), to sequester extracellular calcium, or KN-62 (KN), to inhibit intracellular calmodulin signaling, abrogated the increase in nuclear area and decrease in chromatin condensation observed for non-treated (NT) and vehicle control cells (VH) during 5% cyclic strain routines (**Fig. 4**). Conversely, calcium inhibition only minorly effected the decrease in nuclear area and increase in chromatin condensation during 20% cyclic strain routines. Interestingly, both BAPTA and KN-62 treatments interfered with the slight decrease in nuclear area and increase in chromatin condensation observed during static magnetic field-only routines (MAG).

**Fig. 4.**
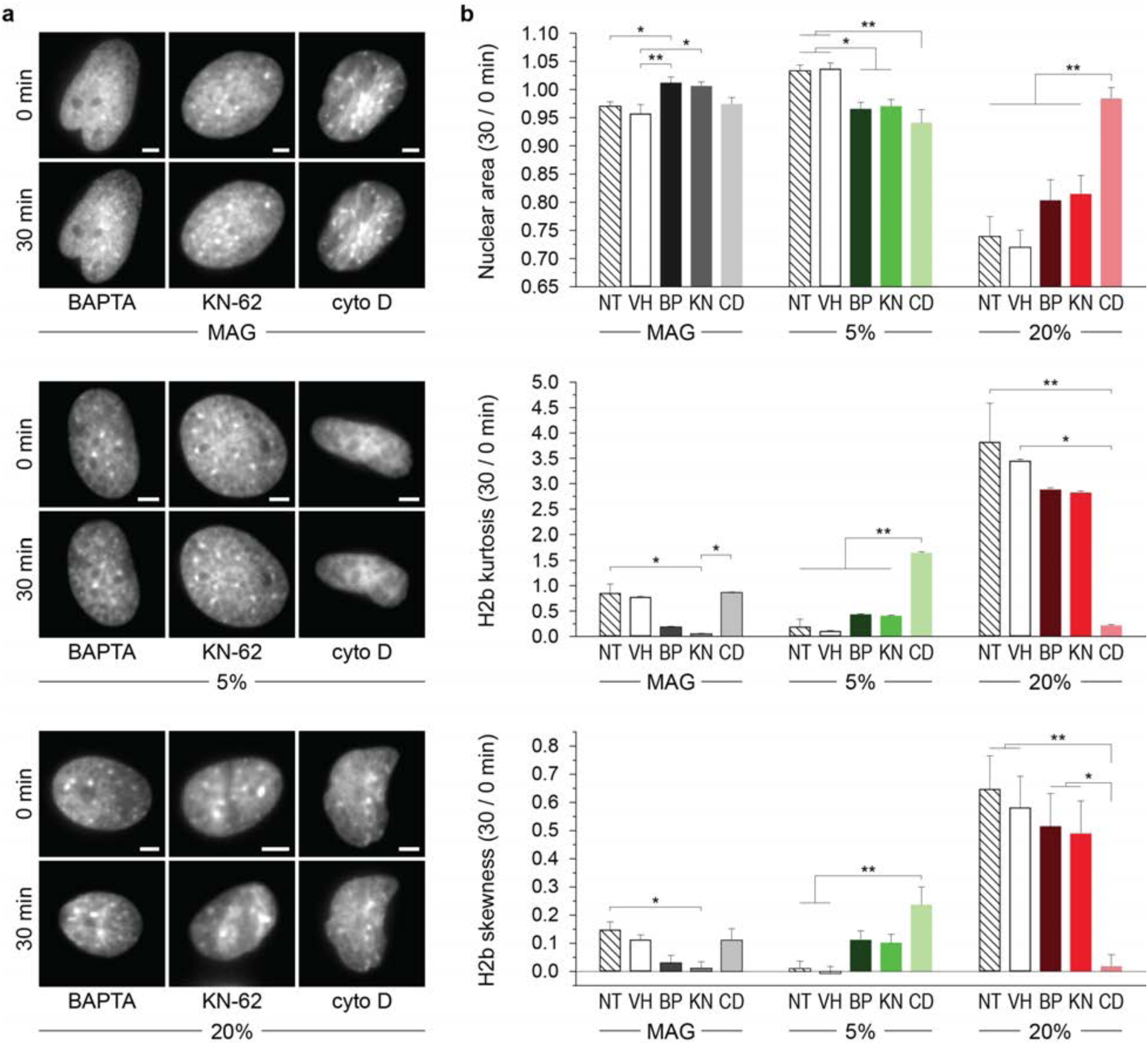
The actin skeleton, but not calcium signaling, is required for nuclear responses to high strain cyclic stretch. Mouse embryonic skin fibroblasts were treated with BAPTA, KN-62 or cyto D before being exposed to 30 min of sigmoidal stretch with peak strains of 5% or 20%, or 0% under the influence of the magnetic field alone (MAG), during which image stacks of nuclei were recorded. **a**) Images of nuclei recorded via H2b-eGFP before (0 min) or after (30 min) cyclic stretch routines; scales=5 µm. **b**) Cyto D treatment (CD) inhibited changes in nuclear area and chromatin condensation in response to 5% and 20% cyclic stretch compared to non-treated (NT) or vehicle (VH) control cells. BAPTA (BP) or KN-62 (KN) treatment abrogated the increase in nuclear area and decrease in chromatin condensation after 5% cyclic stretch but had no effect after 20% cyclic stretch; NT control same as Fig. 3; SEM; n≥15 from 3 exp.; ANOVA: * p<0.05, ** p<0.01.

Further, the actin skeleton has been shown to be crucial for mechanosensitive signaling and is thought to be an important structure for forwarding mechanical cues to the nucleus (18, 21–23, 26). Treatment of MSFs with the actin depolymerization drug cytochalasin D (cyto D, CD) also abrogated the increase in nuclear area in response to 5% cyclic strain and distinctly increased chromatin condensation (**Fig. 4**). Interestingly, cyto D treatment inhibited nuclear shrinkage and chromatin condensation during 20% strain routines. In contrast to calcium inhibition, cyto D treatment showed no effect on MSF nuclei during static magnetic field-only routines. Together, this data suggested that calcium signaling plays a role during low strain, but not high strain stimulation, while an intact actin skeleton was crucial for nuclear responses to any magnitude of stretch.

### Actin Depolymerization Increases DNA Damage After Low and High Cyclic Strain

The calcium-independent shrinkage of nuclear areas suggested that there is a different mechanism for cell behavior under high strain compared to low strain. Cyclic stretch has been shown to cause DNA damage (36, 37). To test whether actin-mediated nuclear shrinkage under high loads might be a protective mechanism to prevent DNA damage, we stained MSFs for serine-139 phosphorylated H2a.x (γH2a.x), to indicate DNA double strand breaks (38), and filamentous actin (F-actin) after 30 min of cyclic stretch. F-actin intensities above the nucleus (perinuclear F-actin) increased with strain magnitudes, being only slightly higher after 5% and twice as high after 20% cyclic strain compared to unstretched (0%) and magnetic field-only control cells (MAG, **Fig. 5a, b**). The number of γH2a.x foci per nucleus also increased with increasing levels of strain. Interestingly, DNA damage was as high under static conditions (0%, MAG) as after 20% cyclic stretch.

**Fig. 5.**
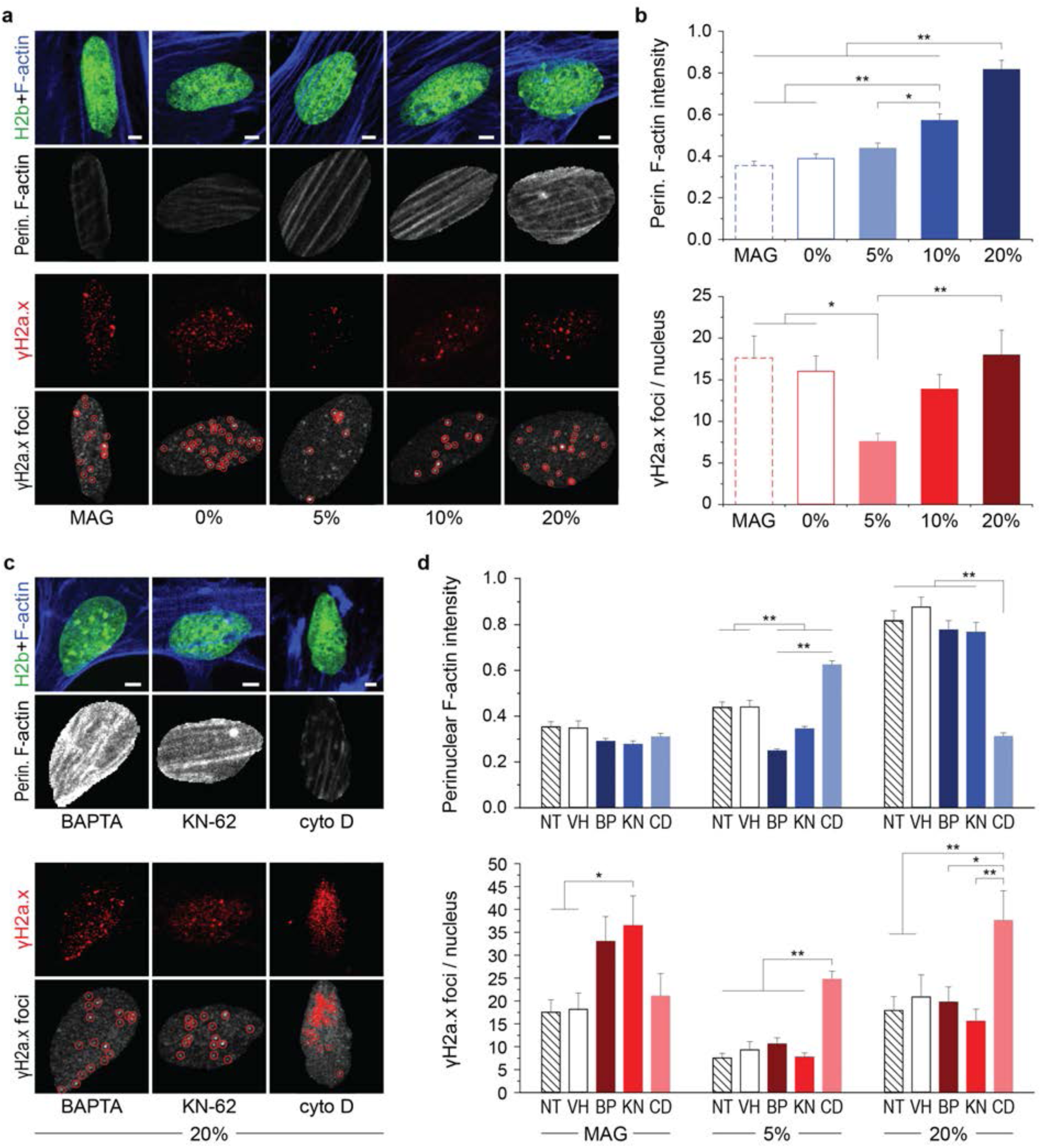
Perinuclear F-actin increases with strain magnitude, and actin depolymerization leads to increased DNA damage after low and high strain cyclic stretch. Mouse embryonic skin fibroblasts were exposed to 30 min sigmoidal stretch routines with peak strains of 0%, 5% or 20%, or under the influence of the magnetic field alone (MAG), after which cells were stained for γH2a.x, as an indicator of DNA double strand breaks, and F-actin. **a**) Stained images of nuclei after stretch routines. A custom MATLAB code was used to analyze perinuclear F-actin intensities, using H2b-eGFP as a mask, and to identify γH2a.x foci as indicated by red circles. **b**) Perinuclear F-actin intensities and number of γH2a.x foci increased with strain magnitude; however, highest levels of DNA damage were observed for static control cells. **c, d**) MSFs were treated with BAPTA (BP), KN-62 (KN) or cyto D (CD) prior to stretch routines. Inhibition of calcium signaling via BAPTA and KN-62 treatment abrogated increases in perinuclear F-actin intensities in response to 5%, but not 20%, cyclic stretch. Actin depolymerization altered F-actin intensities and showed increased number of foci after both 5% and 20% cyclic stretch, while DNA damage was only increased for static magnetic-field only control cells after inhibition of calcium signaling; see Fig. S4a for MAG and 5% images; SEM; n≥150 from 3 exp.; ANOVA: * p<0.05, ** p<0.01; all scales=5 µm.

Similar to nuclear responses, inhibition of calcium signaling via BAPTA or KN-62 abrogated the increase in perinuclear F-actin intensities after 5% cyclic stretch, but not after 20% cyclic stretch, compared to non-treated (NT) or vehicle (VH) control cells (**Fig. 5c, d, Fig. S4a**). Inhibition of calcium signaling increased DNA damage in static magnetic field-only controls but showed no difference for cells after 5% or 20% cyclic stretch. In contrast, actin depolymerization via cyto D treatment distinctly increased the number of γH2a.x foci per nucleus after 5% and 20% stretch routines but had no effect on cells exposed to the magnetic field alone. Cyto D treatment also showed no effect on perinuclear F-actin intensities after magnetic field-only routines. The increase in F-actin intensities after 20% cyclic stretch was abrogated in cyto D treated MSFs, as would be expected after actin depolymerization. However, F-actin intensities were increased after 5% cyclic stretch in cyto D treated MSFs. Judging from the stained images, this was due to an accumulation of small actin filaments at the nuclear periphery (**Fig. S4a**). Further, imaging of nuclei in relaxed or stretched conditions after 30 min of 20% cyclic strain showed that strains transferred to nuclei were higher in cyto D treated cells compared to non-treated cells (**Fig. S4b**). Overall, this data showed that perinuclear F-actin increased with increasing levels of strain and actin depolymerization resulted in elevated occurrences of double strand breaks during high and low cyclic stretch, suggesting that the actin skeleton might play a protective role for the nucleus during high loads.

### Live Imaging of Actin Dynamics Revealed Opposing Patterns of Reorganization During Low and High Cyclic Strain

Inhibition of actin polymerization abrogated nuclear responses to high and low strain routines, and increased DNA damage with increasing levels of strain. To further investigate the role of actin skeleton during stretch-induced changes in cell behavior, we transfected MSFs with a fluorescent F-actin probe (mRuby-Lifeact-7) and acquired image stacks during 30 min of 5% or 20% cyclic stretch followed by 30 min of rest (**Movie 5**). Analysis of 2 µm thick profile line projections along the minor axis (perpendicular to F-actin) of two example cells showed that Lifeact intensities shifted towards the cell border after 30 min of 5% stretch, while they shifted towards the nucleus after 20% (**Fig. 6a, b**). To verify these findings in different cells, profile line projections were grouped into bins along their relative distance to the nuclear center, with bins 1-5 representing Lifeact intensities from the nuclear center to the nuclear periphery and bins 6-10 representing intensities from the cytoplasmic site of the nuclear border to the cells border. After 30 min of cyclic stretch, Lifeact intensities were elevated above the nuclear interior (bin 1-4) in cells exposed to both low (5%) and high (20%) levels of strain (**Fig. 6c**). However, Lifeact intensities were decreased at the nuclear border (bin 6) and increased towards the cell border (bin 10) in cells after 5% of cyclic stretch, while intensities were increased at the nuclear border and decreased at the cell border after 20% of cyclic stretch.

**Fig. 6.**
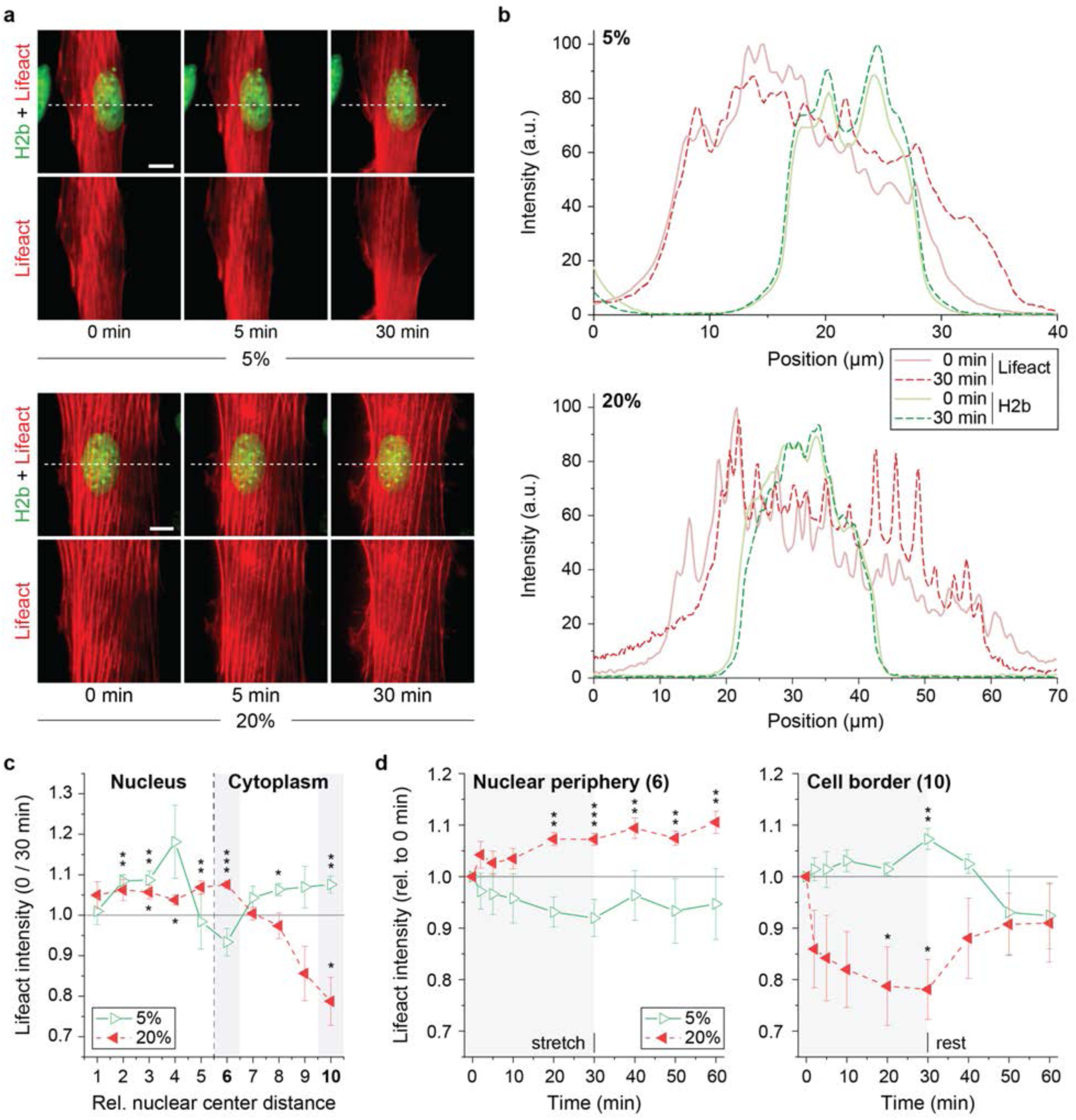
Lifeact imaging reveals opposing changes of actin reorganization at the cell and nuclear border during low strain and high strain cyclic stretch. Mouse embryonic skin fibroblasts were transfected with mRuby-Lifeact-7. The day after, cells were exposed to 30 min of 5% or 20% sigmoidal stretch, followed by 30 min of no stimulation (rest), during which image stacks were recorded. **a**) Images of actin (Lifeact) or nuclei (H2b) recorded during the stretch routine. White doted lines represent the center location of profile lines in (b); scales=10 µm. **b**) Projections of actin (Lifeact) and nuclear (H2b) profile lines, as indicated in (a), of two example cells before (0 min) or after (30 min) exposure to either 5% or 20% cyclic stretch. **c**) Changes in Lifeact intensities after 30 min of stretch were binned into relative location to compare changes in different cells: Bins 1-5 represent intensities from the nuclear center to the inner nuclear border and 6-10 from the nuclear periphery to the cell border. Lifeact intensities shifted from the cell border to the nuclear periphery after 20% cyclic stretch while this trend was inversed after 5% cyclic stretch; SEM; n=6; T-test (vs. 0 min): * p<0.05, ** p<0.01, *** p<0.001. **d**) Changes in Lifeact intensities at the nuclear periphery (bin 6) and cell border (bin 10) over time. Actin reorganization shows dynamics similar to that of nuclear responses; SEM; n=6; T-test (vs. 0 min): * p<0.05, ** p<0.01, *** p<0.001.

Analysis of Lifeact intensities over time showed a steady decrease or incline of intensities at the nuclear border during the 30 min of stretch in response to low or high strain routines, respectively (**Fig. 6c**). Intensities stayed declined or elevated during rest at the nuclear border. At the cell border, Lifeact intensities declined rapidly in response to 20% strain, continued decreasing during the stretch period and raised again during the 30 min of rest, albeit staying lower compared to 0 min. This change in Lifeact intensity was noticeably similar to the dynamics of nuclear area shrinkage after high strain cyclic stretch, suggesting that F-actin rearrangement might mediate the nuclear response through physical interaction. During 5% cyclic stretch, Lifeact intensities slowly increased at the cell border and declined to reduced levels of intensity, compared to 0 min, during rest. In addition, we observed acute nuclear collapse and actin filament disruption in some cells that were exposed to 20% cyclic stretch (**Fig. S5**). These observations were more pronounces after the rest period, indicating that the cell died through either necrosis or apoptosis. This supported the notion that elevated strains are a challenge for cell survival. Taken together, we observed that F-actin also showed opposing patterns of reorganization for different levels of strains, shifting away from the nucleus during low and towards the nucleus during high strain routines. Further, the dynamics of F-actin rearrangements matched that of nuclear responses. Further considering the results from the DNA damage analysis, this suggested that F-actin might mediate the nuclear shrinkage during high loads through physical confinement to provide protection against stretch-induced DNA damage.

## DISCUSSION

In this study, we present a new TENSCell device concept to image cells at high magnification during mechanical stimulation. The device was constructed using mainly 3D printed parts and operated through electromagnetic force. This combination made the device easy to replicate, through the means of 3D printing, while also allowing for the precise control of strains. Another benefit of electromagnets is the simple construction of control circuit and PC-integration, which further adds to the user friendliness. 3D printing also enables the adjustment of the design to fit different microscope stages.

To test this device, we investigated the response of mouse embryonic skin fibroblasts showed nuclei to cyclic stretch. Unexpectedly, we observed opposing responses to sigmoidal strain routines with low or high amplitude, as nuclear areas increased and chromatin decompacted for low strains and areas decreased and chromatin condensed for high strain. Previous studies have shown that cyclic uniaxial stretch (3-15%) induces chromatin condensation and nuclear elongation in the direction of stretch in mesenchymal stem cells (MSC) (17, 18). In another study, cyclic force application using a magnetic needle resulted in chromatin decondensation in HeLa cells (23). All groups reported changes in chromatin compaction to be very rapid upon stimulation, similar to our findings here. However, to our knowledge, no study has reported dichotomous mechanosensitive behavior of nuclei. Pioneering studies in mechanosensation have shown that MSC differentiate in accordance with the stiffness of their environment (1) (becoming osteogenic on stiff, myogenic on medium, and neurogenic on soft substrates) which indicated that cells differentiate between intensities of mechanical cues. Still, the underlying mechanosensitive mechanisms, and whether differences in strain magnitudes are processed through either one pathway in a dose-response manner or are the result of different pathways interacting, is not clear. Here we showed that nuclear responses to low strain cyclic stretch were dependent on calcium signaling, while high strain responses were not, which provided evidence that different pathways are involved in strain magnitude processing.

Based on our observation of divergent nuclear behavior, it could be hypothesized that different magnitudes of strain represent different modes of operation that are associated with specific challenges for the cell. More specifically, low strain cyclic stretch might mimic baseline physical activity and drive regular cell activity or maintenance. In turn, high loads might be associated with extreme activity or trauma and trigger a protective mechanism instead. Increased mechanical stress been shown to cause DNA damage (36, 37) and trigger apoptosis (19, 37). The observed rapid nuclear contraction during high strain loads might therefore be a mechanism to protect from strain-induced DNA damage. In this study, we also observed an increase of DNA double strand breaks with increased levels of strains. Interestingly, the number of double strand break foci was as high in unstretched control cells as after high levels (20%) of cyclic stretch, suggesting that skin fibroblasts cells perform better under dynamic compared to static conditions. It is likely that skin fibroblasts, and other types of cells, have an optimal performance under low mechanical stimulation since that reflects the conditions they evolved in inside motile organisms. This might have grave implication considering that cells are usually cultured on rigid plastic or glass. Hence, more work needs to be done to validate and understand this phenomenon.

We further observed that the actin skeleton was essential for any nuclear response to stretch. Other studies have reported that stretch induced changes in chromatin organization (18, 23, 26) or gene expression (21, 22) were abrogated after disruption of the actin skeleton. One particular study found that the nuclear shape is controlled by a perinuclear actin cap above and around the nucleus (39). Here, we observed that perinuclear F-actin increased with strain magnitudes and would therefore not explain the dichotomous nuclear behavior observed for different strains. However, closer investigation of dynamic F-actin reorganization in live cells showed that F-actin shifted away from the nuclear periphery to the cell border during low strain cyclic stretch and *vice versa* during high strain cyclic stretch. The dynamic change in actin reorganization was also rapid, particularly during high strain routines, and resembled that of nuclear responses. This suggested an interplay between nucleus and cytoskeleton. Moreover, the occurrence of double strand breaks was increased during stretch after actin skeleton disruption. From these results, one could hypothesize that mechanical confinement of the nucleus through F-actin encapsulation might present a mechanism for DNA protection. Further supporting this hypothesis, we observed that strain transferred from the silicone membrane to the nucleus was increased after actin depolymerizing via cyto D after 30 min of cyclic stretch. Nuclear strain transfer prior to cyclic strain application could not be accurately assessed due to the rapid decline in nuclear area of cells held in a stretched position (see methods for details). Therefore, the question whether actin reorganization towards the nuclear periphery reduces nuclear strain transfer remains open. Furthermore, it would be interesting to investigate the role of the LINC complex for the interplay between cytoskeleton and nucleus, since LINC complex disruption has been shown to inhibit nuclear responses to mechanical cues (20, 22, 25) and interferes with cytoskeleton dynamics (25, 40).

A disadvantage of the TENSCell device is the potential influence of magnetic fields, used to operate the device, on cell behavior. The rare earth magnet was positioned away from the containment well at the opposite side of the piston and had a magnetic field strength of 7 mT at the location of the well (>5 cm from the pole). The distance of the magnet from the well changed only slightly (<1 cm) during stretch and can be considered static (±0.1 mT). In contrast, the electromagnetic coil produces an oscillating field during cyclic stretch with a maximum field strength of about 25.4 mT at the level of the membrane (**Fig. S6**) for 20% strain routines (1.0 mT for 5% and −9.3 mT at baseline). Studies showed that long-term exposure (1-5 days) to static 6 mT magnetic fields can have a significant but moderate effect on cell survival and cell morphology for some of the cell lines investigated (41, 42). Reviews on the effects of static magnetic fields concluded that effects on cell survival and proliferation were absent or minor regardless of the field strength used (43, 44). However, it should be noted that static magnetic fields can increase the effect of apoptosis-inducing drugs (41–44) which have been related to field-induced changes in calcium uptake (45, 46). This could also pose a possible explanation for the increase in DNA damage in BAPTA or KN-62 treated cells observed in this study. Investigation of the effect of oscillating electromagnetic fields (1-20 mT) showed moderate effects on cancer proliferation over the duration of 5 or 7 days (47, 48). These studies used 50 Hz oscillations, which are considered low frequency but are distinctly higher than the 1 Hz used in this study. Research in chick embryos showed that oscillating fields influenced development only upwards of 16 Hz independent of the field strength used (49). Overall, studies on static and oscillatory magnetic fields observed only minor changes in cell behavior after long time (days) exposure. In this study, we found no difference in nuclear responses between unstretched control cells exposed to magnetic field routines or cells in the absence of a magnetic field during the 1 h experimental routine. Special caution should be given for the use of pharmacological agents, especially when they negatively affect cell viability.

## METHODS

### TENSCell Device Fabrication

Our custom-built device was designed to acquire high-magnification images of cells during the application of equiaxial strain while also avoiding the use of expensive materials (**Table S1**) or complex designs. Two types of 3D printers were used to print a majority of the components: Objet30 (Computer Aided Technology) using the material VeroClear (OBJ-04055, Computer Aided Technology) and the uPrint SE Plus (311-20200, Computer Aided Technology, Inc) using ABS+. Designs for the device components were created as CAD files (**Fig. S1a**) using SolidWorks (Dassault Systèmes SolidWorks, v. 2018). All parts were printed at the Integrated Teaching and Learning Laboratory (ITLL) at the University of Colorado Boulder.

The main body consisted of four parts: a deformation ring holder, an electromagnet case, a slider tube and a piston (**Fig. 1a, Fig. S1b**), which were printed using the uPrint SE Plus. The deformation ring holder was designed to fit into the circular notch of the manual stage of a Nikon Eclipse Ti Microscope (by interlocking with two metal wings that otherwise hold the aluminum sample tray) and contained an adapter that encased the microscope objective and held the deformation ring. Within the encasement, the objective had a moving range of approximately 8 mm in each direction. The deformation ring was machined from Delrin® Acetal Resin (8572K27, McMaster-Carr) to provide a friction-reduced interface with the silicone membrane. The slider tube fit tightly into the electromagnetic coil and contained 3 slider rails (NS-01-27, Igus) for friction-reduced vertical movement of the piston, which in turn contained 3 matching slider carriages (NW-02-27, Igus). A rare earth magnet (R3525, SuperMagnetMan) was attached to the top of the piston to transmit force from the electromagnet below.

The stretch chamber consisted of three parts: a main chamber, a membrane ring and holding clips (**Fig. S1c**), which were printed using the Objet30. The stretch chamber was assembled by placing a 60×60 mm silicon elastomer membrane (gloss 0.005”, Specialty Manufacturing Inc.) straight onto the elevated inner edge of the main chamber, spanning and fixing the membrane with the membrane ring. The membrane ring was secured laterally with three holding clips. To culture cells, a compliant silicon containment ring was fused to the silicon membrane prior to assembly. Containment rings (d_out_=16 mm, d_in_=11 mm, h=5 mm, A_in_=100 mm^2^) were made from polydimethylsiloxane (PDMS, Sylgard®184, Dow Corning) using a 1:40 mixing ratio to produce soft rings with low mechanical resistance. Circular plastic molds were coated with 3,3,3-Trifluoropropyl-trichlorosilane (452807, Sigma) for 1 h under vacuum after which PDMS was poured into molds and cured over night at 80°C. The contact areas between the silicone membranes and the silicone containment rings were ozone-activated for 60 s via corona arc-discharge (BD-20, Electro-Technic Products Inc.) after which rings were pressed onto the membranes, weighted down with a 100g weigh to maintain close contact and incubated again overnight at 80°C to facilitate bonding. Bonded membranes were sterilized with 70% Ethanol, dried, and stretch chambers were assembled.

### TENSCell Device Control

To operate the stretch device, a simple control circuit was designed in which an Arduino microcontroller (DEV-13975, SparkFun Electronics) modulated the magnitude and direction of a constant 6 A current from a DC power source (9129B, BK Precision) to the coil via a H-bridge (RB-Cyt-132, RobotShop). Two signals from the Arduino to the H-bridge controlled the current flow: A PWM pin (power-wave-modulation) sending low voltage from 0-5 V controlled the current intensity (**Fig. S2a**) and a digital pin (either 0 or 1) controlled the direction of the current to allow lifting of the piston in the relaxed state or attracting the piston downwards to intendent the engaged membrane. Arduino inputs were controlled via MATLAB (Mathworks, v. 2018b) via a USB interface and the Arduino Support from MATLAB package and a custom written code was used to operate the device.

### TENSCell Device Calibration

To measure piston movement and associated membrane indentation in response to electromagnetic fields, a laser distance sensor (Keyence LJ-G5001P) was pointed vertically at the top of the piston and changes in vertical movement were recorded via the Keyence LJ-Navigator software (Keyence, v. 1.7.0.0). Particle tracking (34) was used to determine the amount of strain applied to the membrane in response to electromagnetic fields. For strain measurements, containment wells were coated with 2 µm blue fluorescent beads (F8824, Life Technologies) and images were recorded before and after membrane indentation on an inverted epi-fluorescence microscope (Ti-Eclipse, Nikon) with a 60× water immersion objective (0.26 µm/pix) and an EMCCD camera (iXonEM+, Andor). Bead displacements were determined via the Particle Image Velocimetry (PIV) plugin on ImageJ (NIH, v. 1.50e) and strains were calculated from bead displacements using a custom written MATLAB code (Mathworks, v. 2018b). To determine to baseline Arduino input voltage to keep the piston floating over the deformation ring against its own weight, the piston was placed in a position in which the membrane would not touch the deformation ring and membrane strains during stepwise reduction of the magnetic field were determined. The baseline Arduino input voltage was determined as the input before distinct changes in membrane strain could be observed (**Fig. S2b**). Electromagnetic fields produced by the coil (**Fig. S6**) were modelled using COMSOL (v. 5.2.0.166).

### Mouse Embryonic Skin Fibroblast Isolation, Culture and Pharmacological Treatments

B6.Cg-Tg (HIST1H2BB/EGFP) 1Pa/J mice (Stock No: 006069) were obtained from Jackson Laboratory. All animal procedures were performed following Institutional Animal Care & Use Committee approval. Skin from embryonic mice was harvested 18.5 days post conception. Skin was minced, washed with HBSS and incubated in 35 mm dishes in shallow medium (∼0.5 ml) to avoid floating of the tissue for four days during which fibroblasts extruded from the tissue. After four days, remaining tissue was removed (picked out with a pipette) and extruded cells in the dish were detached using TrypLE (Gibco) and expanded in culture for another two days before being seeded into stretch chambers for experiments. The inner well of assembled stretch chambers was ozone-activated for 30 s via corona arc-discharge (BD-20, Electro-Technic Products Inc.) and coated with 50 µg/ml bovine plasma fibronectin (F1141, Sigma) in a total volume of 250 µl overnight at 37°C. MSFs were seeded into stretch chambers at a density of 80,000 cells/cm^2^ one day before experiments. MSF were extruded and cultured in DMEM (ATCC) containing 10% fetal bovine serum (Gibco), 1% penicillin-streptomycin (Gibco) and 25 mM HEPES (Gibco) at 37°C and 5% CO_2_. To inhibit calcium signaling, MSFs were incubated with 50 μM BAPTA (A4926, Sigma) or 10 μM KN-62 (I2142, Sigma) 1 h prior to cyclic strain experiments. To disrupt actin polymerization, MSFs were incubated with 2 μM cytochalasin D (C8273, Sigma) 30 min prior to experiments. Vehicle controls were incubated with 0.001% DMSO (276855, Sigma) 1 h prior to experiments. To test the use of chromatin dyes, cells were stained with Hoechst (NucBlue™ Live ReadyProbes™, Life Technologies) 30 min prior to experiments.

### Live Imaging and Analysis of Nuclear Behavior During Stretch Routines

The stretch device was mounted on an inverted epi-fluorescence microscope (Ti-Eclipse, Nikon) with a 60× water immersion objective (0.26 µm/pix) and an EMCCD camera (iXonEM+, Andor). Using MATLAB, a sigmoidal control signal (1 Hz) was sent to the Arduino controller, resulting in a sigmoidal stretch routine of the membrane (**Fig. 1c**), with peak strains set to 5% (−0.1 V), 10% (−0.9 V) and 20% (−2.25 V) as determined by the calibration curve. Cells were cyclically stretched for 30 min followed by 30 min of rest (no stretch but the baseline magnetic field keep the piston levitating at 0%) during which 2 µm z-stacks (0.5 µm steps) of nuclei were acquired. Unstretched control cells (0%) were seeded into stretch chambers but kept stationary on the microscope without any magnetic field applied. Unstretched magnetic control cells (MAG) were placed stationary inside the coil without the piston and a magnetic field corresponding to a 20% sigmoidal stretch routine was applied. A custom MATLAB code was written that tracked nuclear outlines and H2b histograms during an image series from which changes in nuclear area, H2b skewness and H2b kurtosis were calculated.

To analyze the amount of strain transferred from the membrane to the nucleus after cyclic stretch routines, image stacks of nuclei were acquired under relaxed or stretched conditions and bulk nuclear strain was calculated from the change in nuclear area using the same MATLAB script as above. Note: nuclear strain transfer could not be accurately analyzed prior to stretch routines due to the fast response in area decline. MATLAB code is available from the corresponding author upon request.

### Perinuclear F-actin and γH2a.x Staining and Analysis

MSFs were fixed in 4% ice-cold PFA for 10min, permeabilized with 1% Triton-X100 in PBS for 15 min and blocked with 10%NGS, 1% BSA in 0.1% PBT (0.1% Tween-20 in PBS) for 60 min. Primary incubation of Phospho-Histone H2A.X Ser139 (Cell Signaling, 9718S) was performed at 4°C overnight in 0.1% PBT containing 1% BSA at 1:400. Secondary incubation of Goat anti-Rabbit IgG-AF546 (Life Technologies) was performed in primary incubation buffer for 45 min at 22°C at a dilution of 1:500. F-actin was counterstained with Phalloidin-CF405 (Biotium) for 30 min at a dilution of 1:30. Staining was performed in containment wells. After staining, membranes were cut out of the containment well, using a circular punch, and mounted cell-side up onto #1.0 glass slides with ProLong™ Diamond Antifade Mountant (Life Technologies). Image stacks (5 µm, 1 µm step) of multiple nuclei in a 318×318 µm area were acquired on a Nikon A1R confocal microscope using a 40× oil immersion objective (0.31 µm/pix). A custom MATLAB code was written that identified nuclei and determined perinuclear F-actin intensities and γH2a.x foci in projected image stacks. To account for variations in staining and imaging (same imaging settings were used), fluorescence channels were histogram normalized. Perinuclear F-actin intensities were calculated as the sum of normalized Phalloidin intensities within a nuclear border. DNA damage foci were determined by detecting 2D peaks in the normalized γH2a.x channel using the MATLAB script FastPeakFind (v. 1.7) previously programmed by Adi Natan. MATLAB code is available from the corresponding author upon request.

### LifeAct Transfection and Analysis of Actin Dynamics

MSFs were transfected with mRuby-Lifeact-7 using Lipofectamine 3000 (Life Technologies) 18 h after seeding into stretch chambers; mRuby-Lifeact-7 was a gift from Michael Davidson (#54560, Addgene). One day after transfection, 2 µm image stacks (0.5 µm steps) of Actin (Lifeact) and Nuclei (H2b) were acquired during cyclic stretch and the following period of rest. Profile lines were generated with the Plot Profile function in ImageJ (NIH, v. 1.50e) using an 8 pixel (=2.1 µm) thick line. Binned profile analysis was performed using a custom MATLAB code that tracked cell and nuclear outlines during an image series using the Lifeact or H2b fluorescence channel, respectively. Image stacks were rotated to align actin filaments in the horizontal direction and projections of 8 pixel thick vertical profile lines that crossed through the nuclear center were extracted for further analysis. During each time step, profile line positions were fixed with respect to the nuclear center. Using the position of the cell and nuclear boundary, profile lines were binned with 5 bins representing Lifeact intensities from the nuclear center to the inner nuclear border and another 5 bins representing intensities from the nuclear periphery to the cell border. Corresponding bins from each half of the cell were averaged. MATLAB code is available from the corresponding author upon request.

### Statistical Analysis

One-way ANOVA with Tukey’s Honestly Significant Difference post hoc test or two-tailed T-test analysis was performed to evaluate statistical significance using JMP Pro12 software (SAS Institute). Displayed error (SD=standard deviation, SEM=standard error of the mean), number of individual data points (n), number of independent experiments (exp., if different from n) as well as significances and statistical tests that were used are indicated in the figure captions.

## ACKNOWLEDGEMENTS

We want to thank the Integrated Teaching and Learning Laboratory (ITLL) at the University of Colorado Boulder for help and advice with 3D printing. This work was supported in part by NIH grants R01 AR063712 and R21 AR066230, and NSF grant CMMI CAREER 1349735.

## AUTHOR CONTRIBUTIONS

Conceptualization, B.S., C.P.N.; Methodology, B.S., A.K.S, I.N., K.C., C.P.N.; Software, B.S.; Formal Analysis, B.S., S.E.S.; Investigation, B.S., A.K.S and I.N.; Writing – Original Draft, B.S.; Writing – Review & Editing, All authors; Funding Acquisition, C.P.N.; Resources, C.P.N.; Supervision, C.P.N.

## DECLARATION OF INTERESTS

Patent pending, provisional Patent Application No. 62/876,535.

## SUPPLEMENTAL INFORMATION

### SUPPLEMENTAL TABLES

**Table S1.**
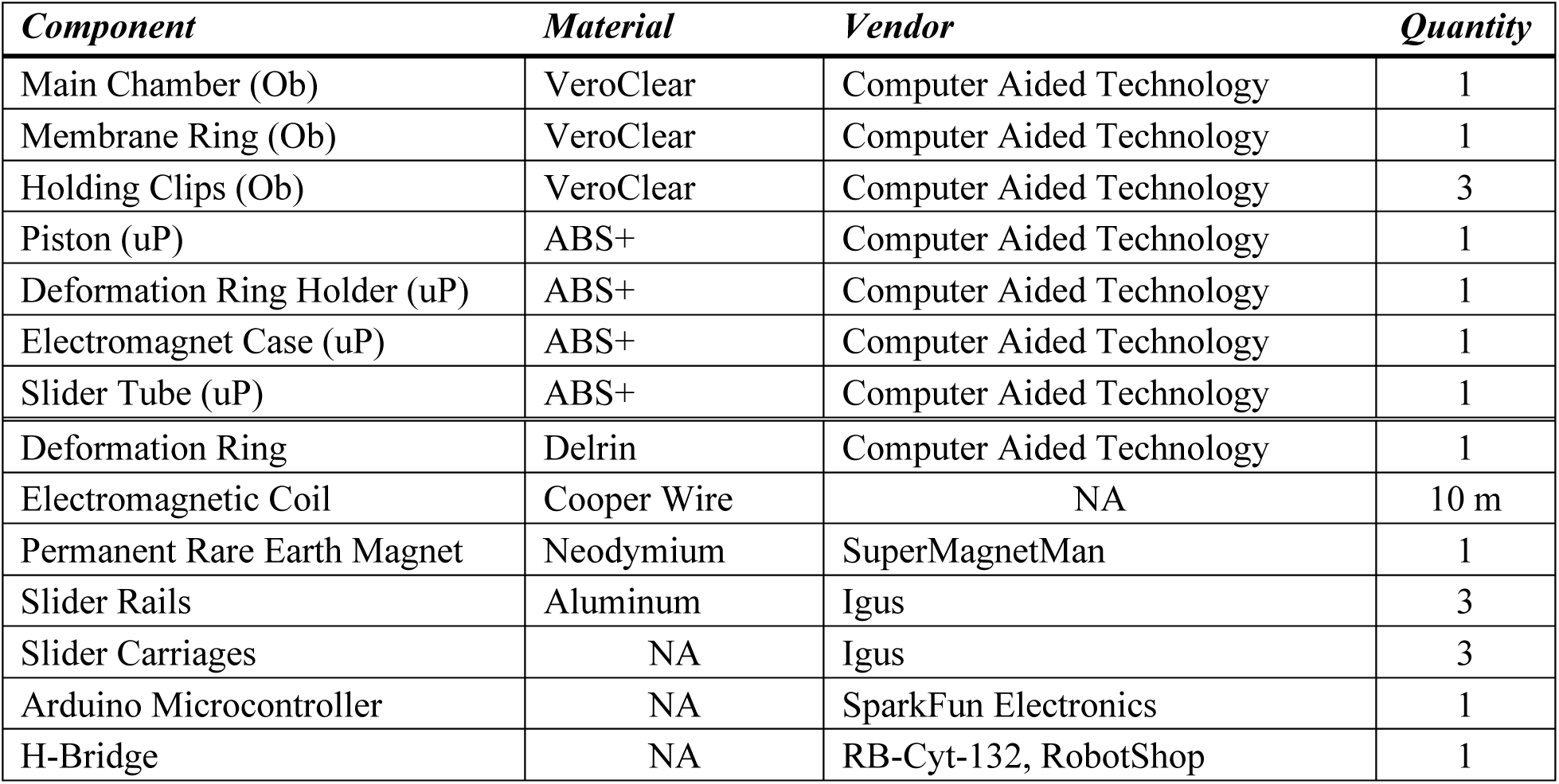
Component list of the TENSCell imaging device. Pricing for the device depends on materials costs and 3D printing, including cost of printing material, density of the print, and facility charges. Parts were printed with the Objet30 3D prints (Ob) or uPrint SE Plus (uP) as indicated.

| *Component* | *Material* | *Vendor* | *Quantity* |
| --- | --- | --- | --- |
| Main Chamber (Ob) | VeroClear | Computer Aided Technology | 1 |
| Membrane Ring (Ob) | VeroClear | Computer Aided Technology | 1 |
| Holding Clips (Ob) | VeroClear | Computer Aided Technology | 3 |
| Piston (uP) | ABS+ | Computer Aided Technology | 1 |
| Deformation Ring Holder (uP) | ABS+ | Computer Aided Technology | 1 |
| Electromagnet Case (uP) | ABS+ | Computer Aided Technology | 1 |
| Slider Tube (uP) | ABS+ | Computer Aided Technology | 1 |
| Deformation Ring | Delrin | Computer Aided Technology | 1 |
| Electromagnetic Coil | Cooper Wire | NA | 10 m |
| Permanent Rare Earth Magnet | Neodymium | SuperMagnetMan | 1 |
| Slider Rails | Aluminum | Igus | 3 |
| Slider Carriages | NA | Igus | 3 |
| Arduino Microcontroller | NA | SparkFun Electronics | 1 |
| H-Bridge | NA | RB-Cyt-132, RobotShop | 1 |

### SUPPLEMENTAL FIGURES

**Fig. S1.**
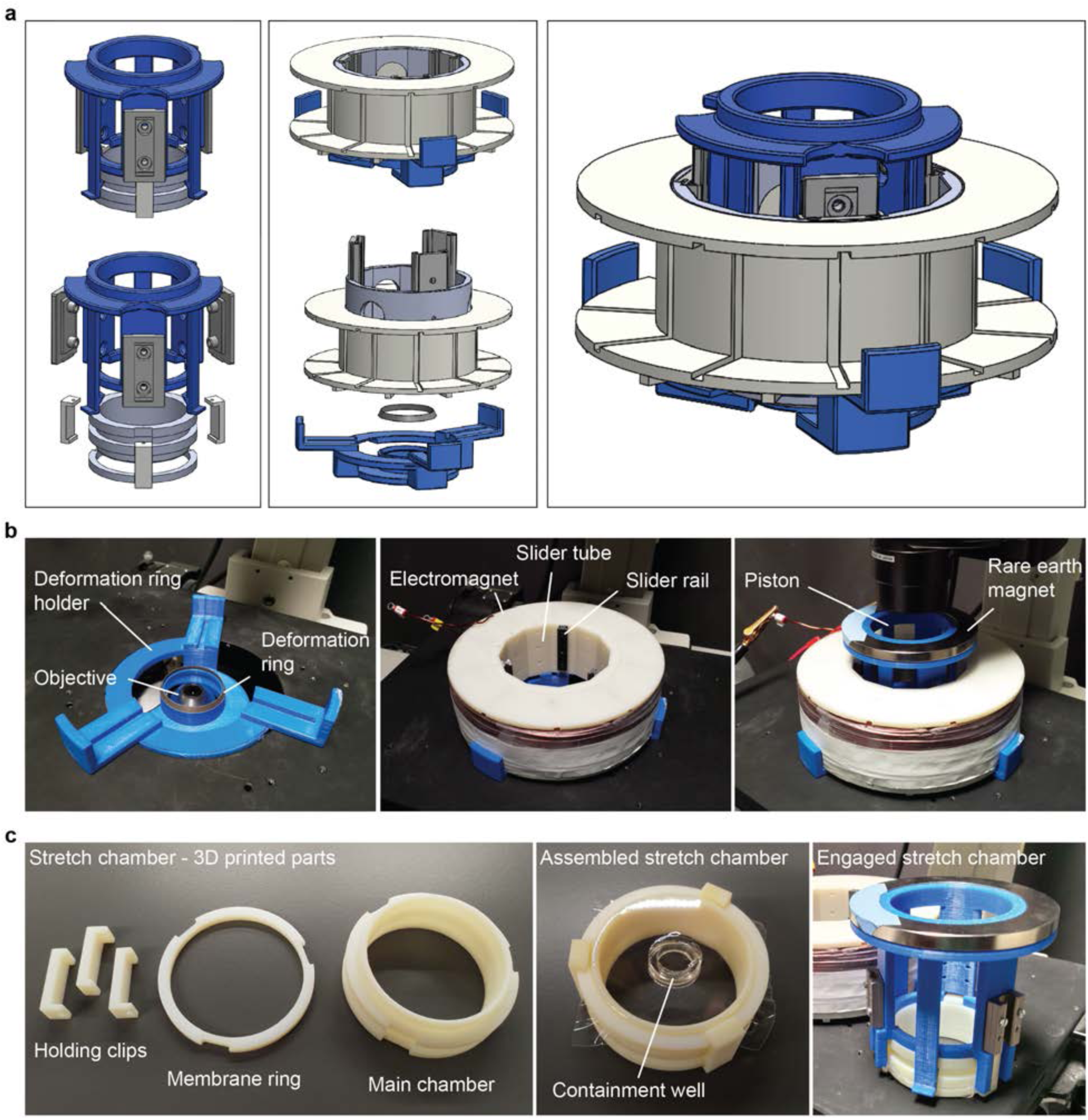
CAD sketches and real images of TENSCell device components and assembly. **a**) SolidWorks CAD drawings of the 3D printed stretch device components shown assembled or as blow-up. Drawings also show slider rails and carriages (in gray), which were purchased. **b**) The deformation ring holder is fixed into the manual stage of a Nikon Eclipse Ti Microscope by interlocking with two metal wings in the circular notch that otherwise holds an aluminum tray. The slider tube pushed tightly into the inner hole of the electromagnetic coil. Together, the slider tube and electromagnetic coil are placed on top of the deformation ring holder. The piston attached to a rare earth magnet is then placed into the slider tube via the friction-reduced slider interface. **c**) The stretch chamber consisted of three parts which after assembly would fix a thin silicone membrane in place. A compliant silicon containment ring was fused to the silicon membrane prior to assembly to culture cells. The assembled stretch chamber can then be engaged into the piston and is held securely in place through bendable side pins that lock into the side grooves of the main chamber.

**Fig. S2.**
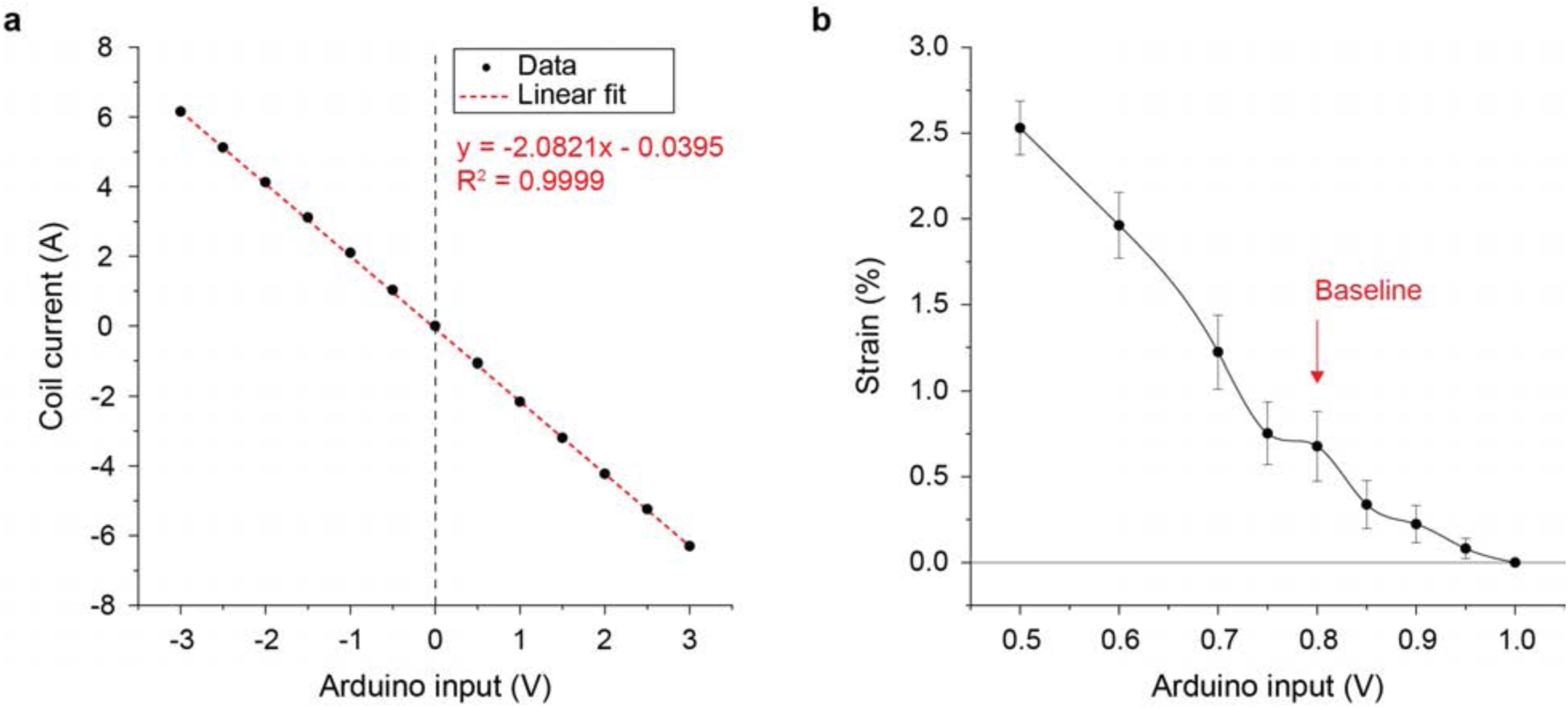
Current control via Arduino input voltages and determination of baseline Arduino input voltage. **a**) Correspondence of Arduino input voltages to the amount of current received by the electromagnet coil was measured using a multimeter. **b**) The baseline Arduino input voltage is the input at which the membrane contacts the deformation ring without indention. Membrane containment wells were coated with 2 µm sized fluorescent beads and membrane strains were calculated using particle tracking. To determine the baseline, the membrane was suspended above the deformation ring and the Arduino input voltage was stepwise lowered (+1.0 to +0.5 V) while measuring the strain applied to the membrane via bead displacements. The baseline Arduino input voltage was determined to be +0.8 V as the input before distinct increases in membrane strain could be measured.

**Fig. S3.**
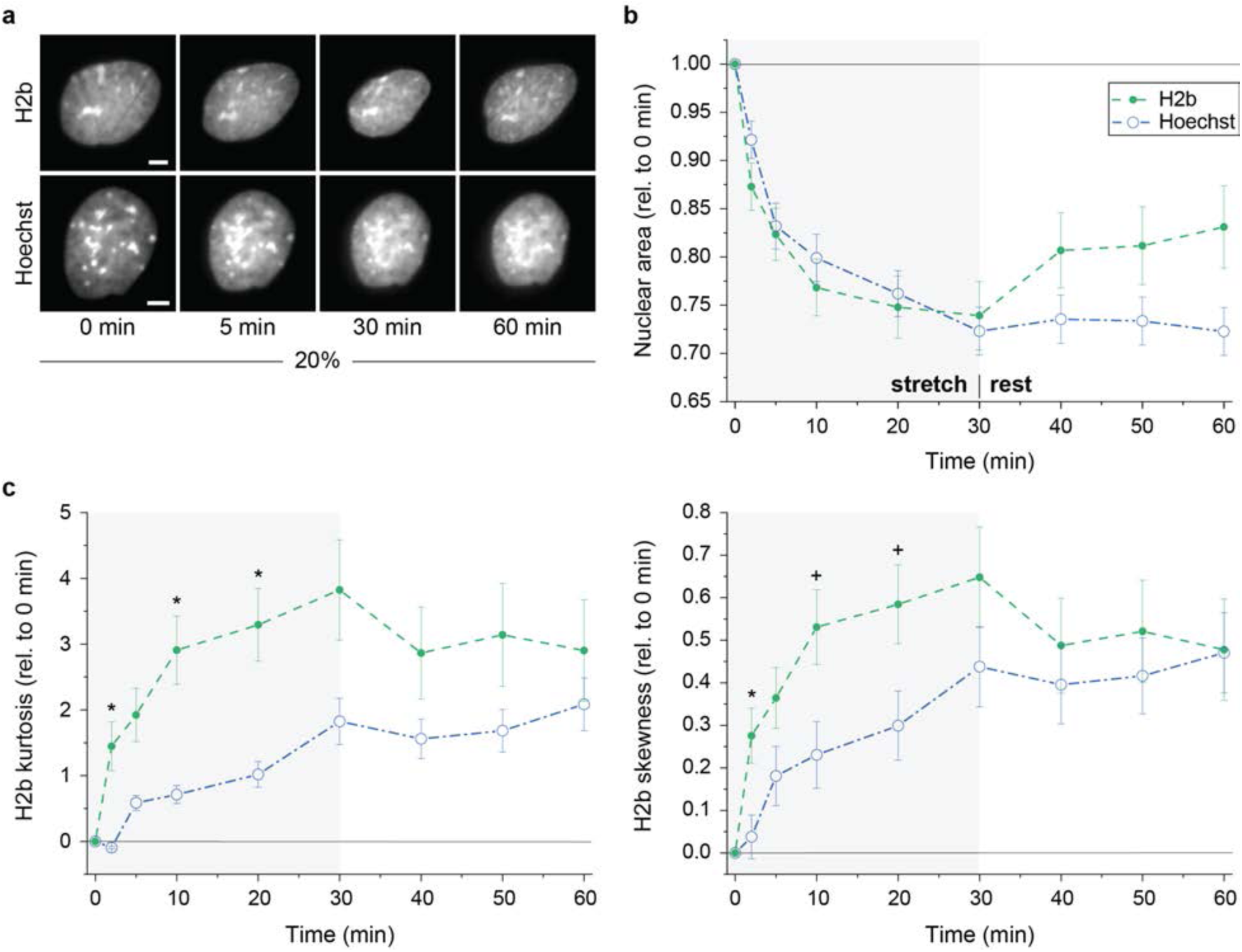
Use of chromatin dyes changes nuclear response to high strain cyclic stretch in MSFs. Mouse embryonic skin fibroblasts were exposed to 20% cyclic strain for 30 min followed by 30 min of no stimulation (rest) during which image stacks of nuclei were recorded. To test the influence of chromatin dyes, cells were stained with Hoechst 33342 prior to experiments. **a**) Images of nuclei recorded via H2b-eGFP or Hoechst staining; scales=5 µm. **b, c**) Relative changes in nuclear area (relative to 0 min) and differences in H2b histogram kurtosis and skewness (compared to 0 min) during stretch routines between DAPI stained or unstained control cells (H2b). Hoechst stained cells showed reduced chromatin condensation during stretch and no regain in nuclear areas during rest; SEM; H2b same as Fig. 3; Hoechst n=10 from 2 exp.; T-test: + p<0.1, * p<0.05.

**Fig. S4.**
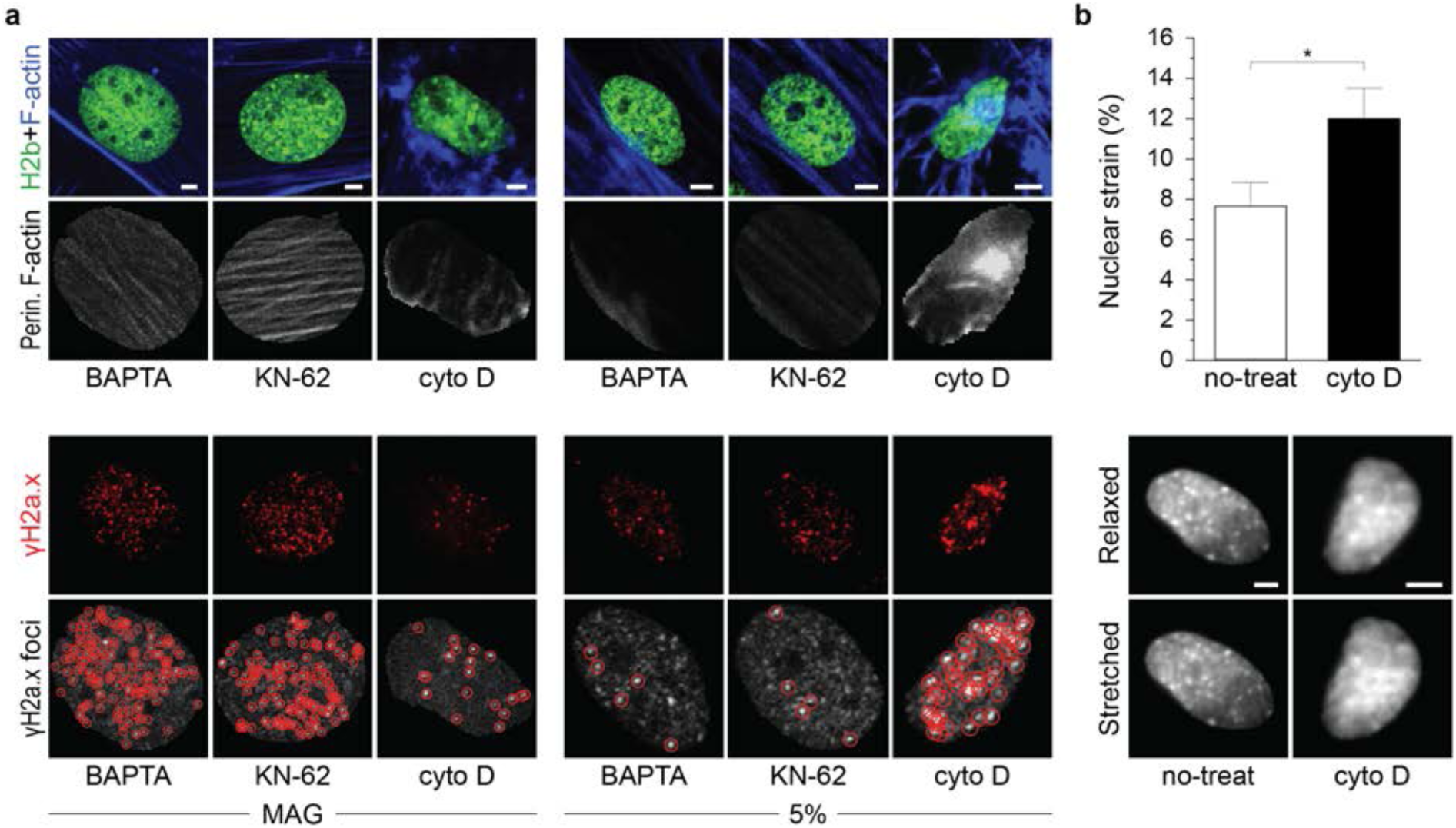
Cyto D treated MSFs show higher nuclear strain transfer during stretch. Mouse embryonic skin fibroblasts were exposed to 30 min sigmoidal stretch routines after which cells were stained for γH2a.x, indicating DNA double strand breaks, and F-actin. **a**) Additional immunostaining data for 5% cyclic stretch and magnetic field only (MAG) routines corresponding the Fig. 5. **b**) After 20% cyclic stretch, images of nuclei were recorded under relaxed or stretched conditions to determine strain transfer from the membrane to nuclei. Cells treated with cyto D prior to experiments showed higher nuclear strains compared to untreated (no-treat) control cells; SEM; n=28 from 4 exp.; T-test: * p<0.05; all scales=5 µm.

**Fig. S5.**
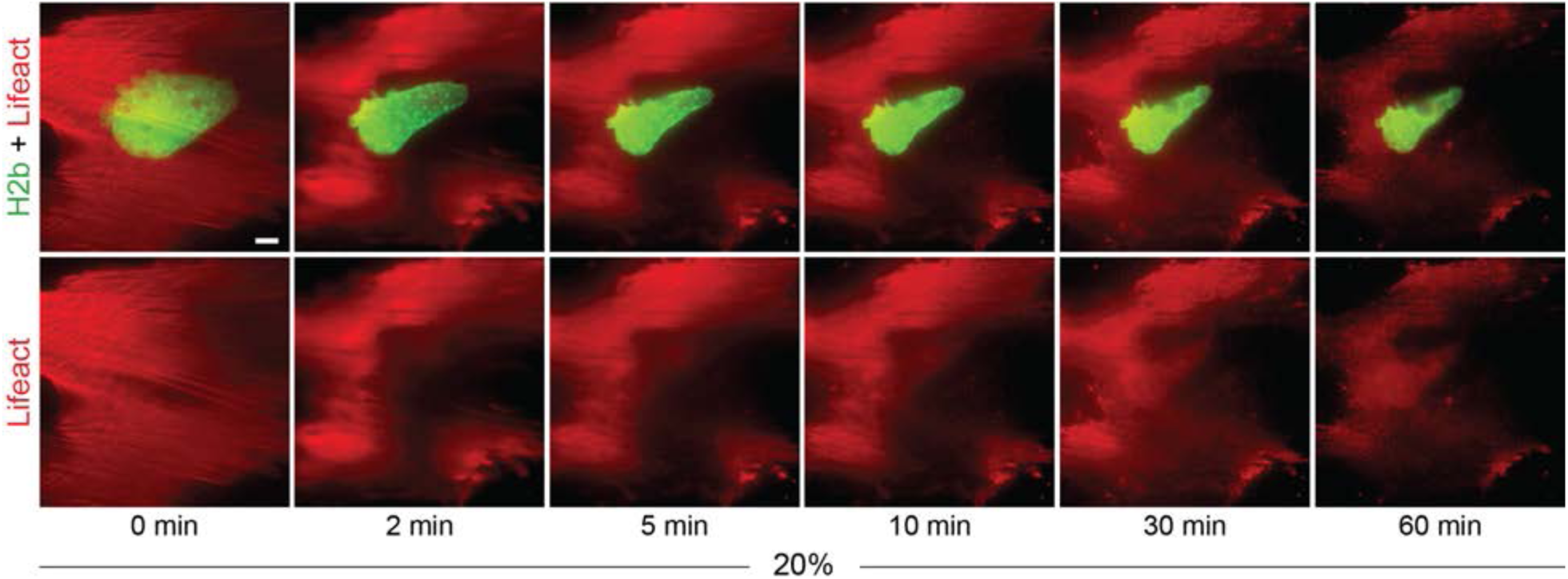
Lifeact imaging of a cell undergoing cell death during high strain cyclic stretch. Mouse embryonic skin fibroblasts were transfected with mRuby-Lifeact-7 and, 24 h after, cells were exposed to 30 min of 20% sigmoidal stretch, followed by 30 min of no stimulation (rest). Image stacks of actin (Lifeact) or nuclei (H2b) were recorded during the stretch routine. Image series shows a cell undergoing apoptosis shortly after start of the high strain routine; scale=10 µm.

**Fig. S6.**
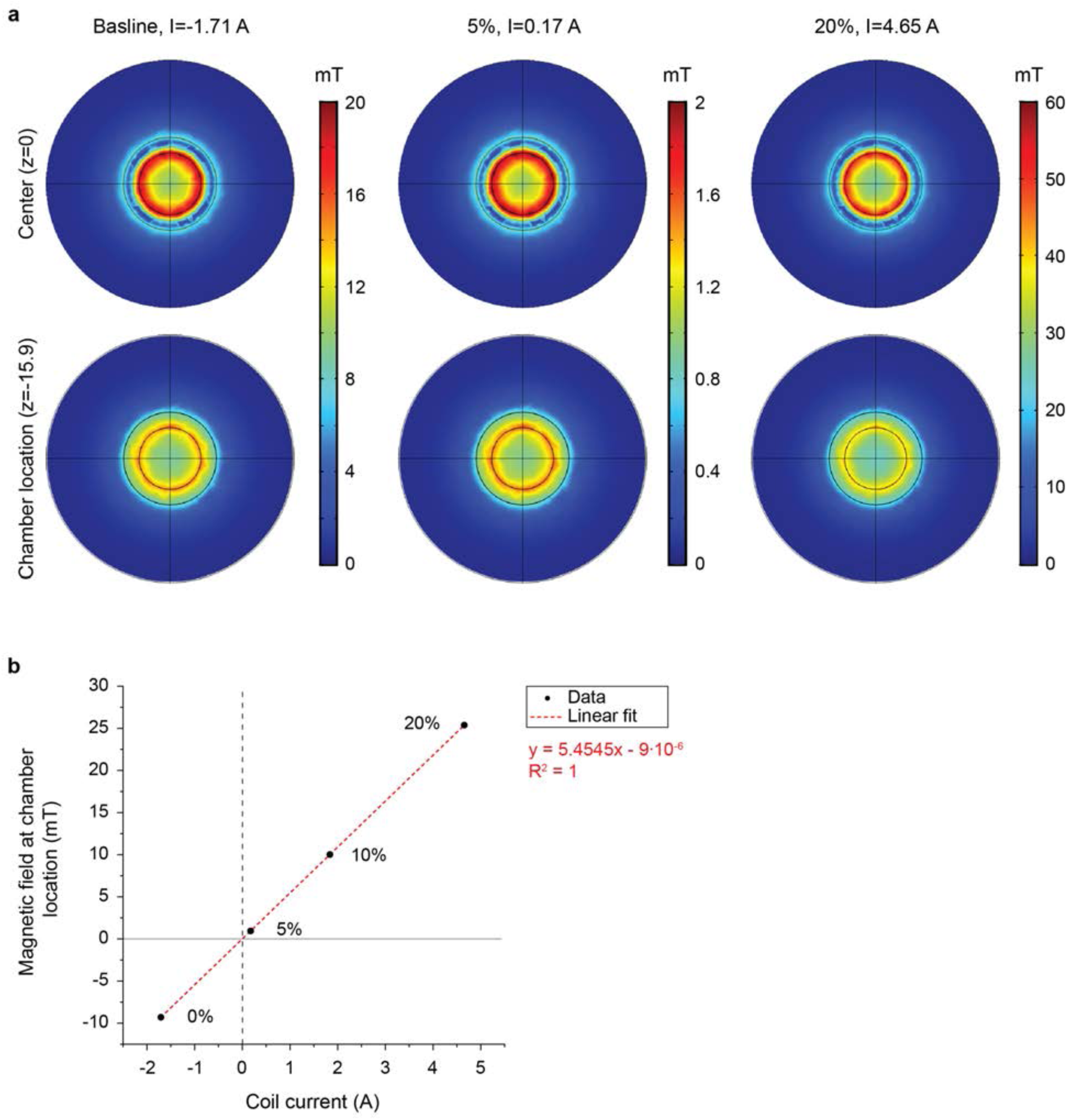
COMSOL-modeling of electromagnetic fields. **a**) Top view of magnetic fields produced by the electromagnetic coil during peak strains corresponding to 5% or 20% strain routines, or during rest (baseline). Two different z-planes are presented: at the center of the coil (*z*=0 mm) and at the location of the membrane close to the bottom of the coil (*z*=15.9 mm). Amount of current used to produce magnetic fields are indicated at the top. **b**) Magnetic fields at chamber location (*z*=15.9 mm) for different currents applied to the electromagnetic coil. Corresponding levels of membrane strain are as indicated.

